# Object motion representation in the macaque ventral stream – a gateway to understanding the brain’s intuitive physics engine

**DOI:** 10.1101/2024.02.23.581841

**Authors:** Hamidreza Ramezanpour, Filip Ilic, Richard P. Wildes, Kohitij Kar

## Abstract

Effective interaction with moving objects and the ability to infer and predict their motion (a core component of “intuitive physics”) is essential for survival in the dynamic world. How does the primate visual system process such stimuli, enabling predictive capabilities for dynamic stimuli statistics like motion velocity and expected trajectories? In this study, we probed brain areas in the ventral visual pathway of rhesus macaques implicated in object recognition (areas V4 and inferior temporal, IT, cortex) to evaluate how they represent object motion speed and direction. We assessed the relationship between the distributed population activity in the ventral stream and two distinct object motion-based behaviors—one reliant on information directly available in videos (speed discrimination) and the other predicated on predictive motion estimates from videos (future event predictions). Further, employing microstimulation strategies, we confirm the causal, functional role of the IT cortex in these behaviors. Our results underscore the need to re-examine the traditional functional segregation of the primate visual cortices into “what” and “where” pathways and provide empirical constraints to model their interaction for a better circuit-level understanding of visual motion and intuitive physics.

## Introduction

Primates live in a dynamic and complex environment in which the integration of various sources of information about objects is essential for survival. Motion and form are two fundamental aspects of intuitive physics for primates, as they enable the recognition of objects in the environment and the prediction of their behavior, ultimately facilitating the primate’s ability to interact with and navigate their surroundings ^1^.

The processing of various object features such as object shape, position, and motion in the visual system has been traditionally discussed in the context of the two-stream hypothesis comprising two anatomically and functionally disparate neural pathways in the primate brain – the dorsal and the ventral stream ^2–4^. Over the last decades, much progress has been made in identifying the brain areas ^5–9^ and the underlying computations ^10–13^ implicated in visual object recognition in the primate ventral visual pathways. On the other hand, mechanisms of visual motion perception have been tied almost exclusively to the dorsal stream ^14–16^. One prominent feature of such studies while probing object recognition is the predominant use of stationary stimuli, and for studies on motion perception, the use of controlled stimuli like random dot kinematograms or moving gratings. Notably, decades of these studies have yielded a significant understanding of static object processing in the ventral stream ^9^ and motion processing (specifically of low-level visual cues like dots and gratings) in the dorsal stream^17^. However, higher-level cognitive functions primarily involve the motion of visual objects, and it is unclear whether inferences made about the neural mechanisms of static object perception and low-level motion cues are sufficient to develop large-scale models of the visual system that can mimic primate brain mechanisms and behaviors in real-world, dynamic tasks ^18,19^.

To represent object motion and support primates’ intuitive physical understanding of the world, it is reasonable to assume that object form and motion information must be integrated at some point (sometimes also considered part of the “binding problem” ^20^). Does the object form information available in the ventral stream combine with motion information extracted from temporal image streams in the dorsal stream? If so, where and how? This integration could occur independently within either stream, downstream of the inferior temporal (IT) cortex, or through a combination of all these pathways. The complexity of this integration makes it challenging to test these theories conclusively. We reasoned that falsifying the independence of the paths could be a promising starting point for exploring the hypotheses further. Therefore, in this study, we exclusively probe the ventral stream during motion-based tasks. Why the ventral stream? While several studies ^21,22^ have highlighted the presence of object form information in the dorsal stream, the richness and specificity of object representation in the ventral stream ^6–8^ and its causal links to object-based behavior ^23^ make it our first circuit of choice. In addition, given that retinotopic maps are present throughout the visual system (including the IT cortex ^24^), information about object position might be present in the activity across most visual areas. Consistent with this notion, a previous study ^25^ has already provided evidence that the IT cortex, classically thought not to be involved in spatial processing, can also represent the object’s location. Hence, it is reasonable to assume that the object position information could be temporally integrated to synthesize motion velocity signals available within the IT cortex or downstream regions like the caudate ^26^ and the prefrontal cortex ^27^. Furthermore, electrophysiological studies have also shown that some IT neurons respond to gratings solely defined by motion ^28^, as well as more complex forms of motion present in biological actions ^29,30^. Also, it has been shown that IT neurons respond more strongly to moving shapes that are degraded than clear ones ^31^. Despite these preliminary findings, how such information relates specifically to object motion speed and direction and links to behavioral measures remains poorly understood – and hence is a significant focus of this study.

This study employed a multifaceted approach to investigate the ventral stream’s role in the primate visual system’s ability to process and predict dynamic stimuli. We began by mapping the neural responses to static and dynamic scenes, discerning the nature of object motion speed and direction representation (in relation to object identity representation) within areas V4 and IT. By formulating temporal and instantaneous decoding strategies (linking hypotheses), we assessed the IT cortex’s role in velocity discrimination and predictive tasks that probe the animals’ capability to develop intuitions about the physical properties of objects, like their motion. Our results demonstrate the causal role of the ventral stream during dynamic, object motion-based tasks and challenge the traditional segregation of the “what” and “where” pathways in the visual cortex, suggesting a more integrated approach to understanding motion processing and predictive capabilities. Through these experimental tactics and inferences, our study contributes to a more nuanced model of visual motion and intuitive physics at the circuit level within the primate brain.

## Results

As outlined above, a critical premise of our study is that the distributed population activity across the macaque IT cortex provides accurate estimates of not only an object’s identity ^7,8^ but also its position ^25^. For instance, for an image of a plane (shown in **Figure 1A, left panel**), three latent variables (out of many) could be reported – this object is a plane at the horizontal and vertical positions of x_1_ and y_1_, respectively. **Figure 1A (right panel)** demonstrates one possible way (as inferred in previous studies ^25^) to decode these latent variables from the IT population activity (by linear weighted summations of the population response). Therefore, we start by replicating object position estimation from IT population activity. We performed large-scale neural recordings across the IT cortex in 3 macaques (total 600 sites) while they passively viewed images presented foveally within 8 deg of visual angle (640 images, eight objects, 80 images per object; *see Methods*) for 100 ms each with multiple repetitions. We then trained regularized linear regression models (cross-validated) to map the population IT activity onto the object’s horizontal (x) and vertical (y) positions. Consistent with previous results, we observed that these regression models could use IT activity from held-out images to predict the corresponding spatial positions of objects with high fidelity (Pearson R = 0.73 for x-position and 0.77 for y-position with 600 neural sites, **Figure 1B**). We also observed an increase in the precision of the position estimates with a larger sample size of neural sites. This scaling of accuracy with the number of sampled multiunit sites was consistent with previous reports (as indicated by the dashed lines representing reported correlation strengths ^25^, **Figure 1B**), bolstering our confidence in the replicability and reliability of our approach.

**Figure 1.**
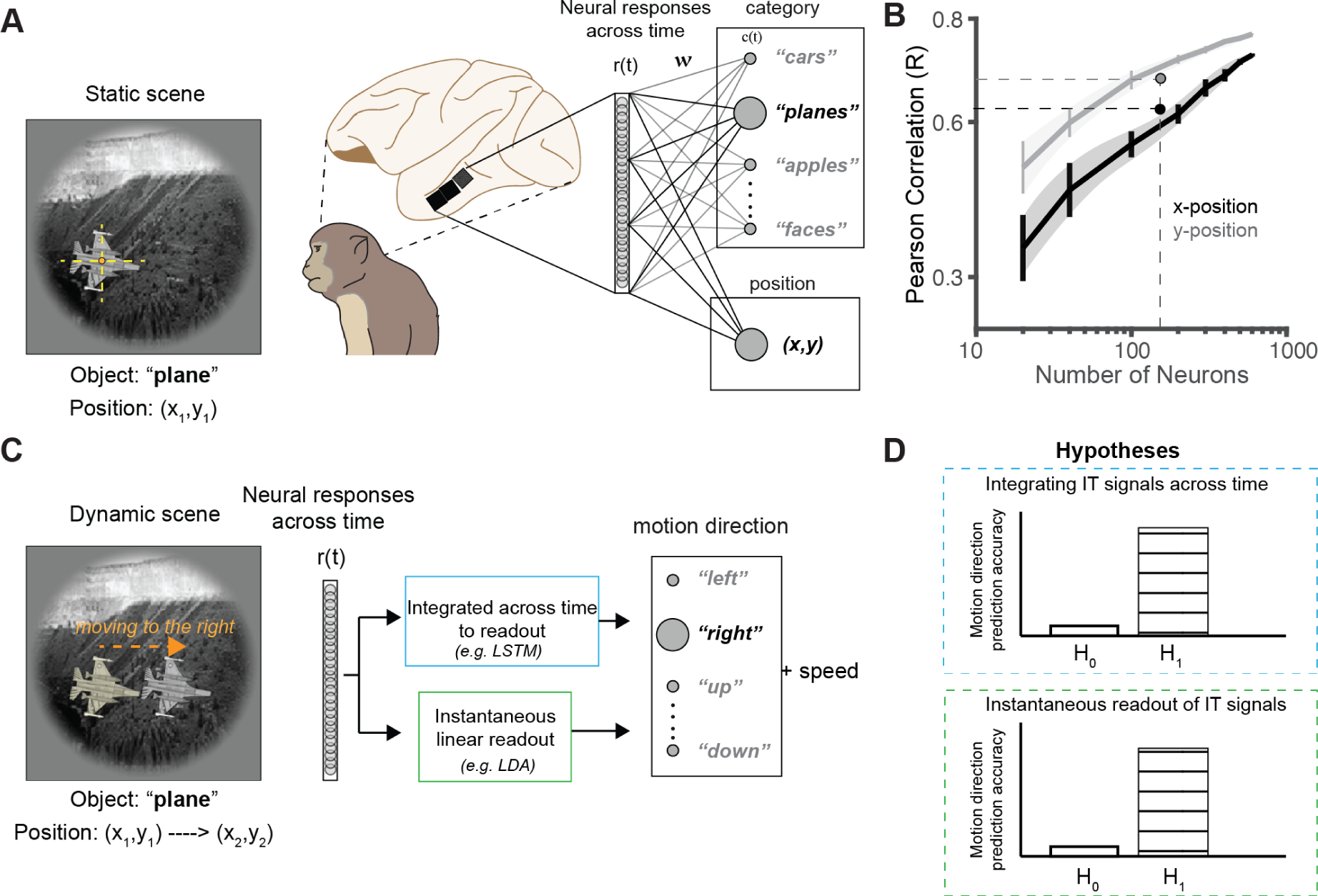
Object position estimates and hypotheses for decoding motion variables from the IT population activity. A. Illustration of the neural response to a static scene in the primate visual system. The image depicts a plane at coordinates (x_1_, y_1_) and showcases how the neural activations within the IT cortex can be mapped to object categories^8^ and spatial positions^25^ (as downstream linking propositions). **B.** Pearson correlation between the number of neurons and their ability to predict object position. With 600 neural sites, we achieved correlations of 0.73 and 0.77 for the x and y positions, respectively. We observed increasing precision in position estimation with a larger neural population. Dashed lines indicate previous reports of correlation strengths with 168 IT sites ^25^, consistent with our results. **C.** Dynamic scene processing showing the same object, a plane, now moving to the right at a certain speed. The image conveys how neural responses, r(t), evolve over time and can be analyzed through time-integrated models (e.g., LSTM) or instantaneous linear readout methods (e.g., linear discriminant analysis, LDA) to decode motion direction and speed. **D.** Hypothetical outcomes of the two proposed readout strategies, suggesting predictions for motion direction prediction accuracy. H_0_ represents the null hypothesis (no difference), and H_1_ represents the alternative hypothesis (a difference), with the upper panel for integrated signal readouts and the lower panel for instantaneous readouts.

Given these results, we next asked whether IT population activity could also predict aspects of the object’s change in position – motion (e.g., the plane moving to the right at a specific speed, **Figure 1C**). We parameterized object motion in terms of its direction and speed. In the following sections, we address the ability of area V4 and the IT cortex to predict each component in turn. One possibility is that neural activity that contains information about an object’s instantaneous position could be integrated over time to represent an object’s direction and speed. For instance, a long short-term memory (LSTM) network-based decoding strategy^32^ that utilizes the full dynamic signal from the IT cortex could be leveraged to estimate the object’s motion variables. On the other hand, one could also ask whether instantaneous activity in IT could be linearly combined to estimate the motion variables (similar to the position estimates). Two potential outcomes of these decoding strategies are visualized in **Figure 1D**. If we cannot falsify the integrated IT signal readout hypothesis (**Figure 1D**, top panel, H_1_), it would suggest that neurons downstream of the IT cortex (or within the IT cortex) could rely on a temporal accumulation of the IT population responses to decode motion velocity accurately. Conversely, if the instantaneous readout hypothesis cannot be falsified (**Figure 1D**, bottom panel, H_1_), it would suggest that neurons within the IT cortex contain linearly separable motion information that accurately enables instantaneous motion velocity decoding. Alternatively, it is also possible that one could only decode object positions for stationary objects from the IT cortex, and this ability is lost when the object is set in motion (H_0_, **Figure 1D**).

### Cortical response dynamics across the ventral stream can be integrated to predict object motion direction accurately

We first assessed the efficacy of temporally integrative decoding strategies to test how well we can decode object motion direction from the population activity distributed across the primate ventral stream. To this end, we recorded large-scale activity across the V4 and IT cortices in two macaques while they passively fixated on brief videos (600 ms) presented at the central 8 deg (*see Methods*). We employed Long Short-Term Memory (LSTM) networks ^32^ to process the dynamic neural responses from area V4 and the IT cortex. We trained an LSTM network with 200 hidden units (**Figure 2A**, *see Methods*) to integrate neural responses (of 192 IT and V4 sites, respectively) over a specific time window (starting from the video onset) and predict the motion direction of the visual objects (400 videos, ten objects, 40 videos per object, eight evenly distributed motion directions). Both V4 and IT responses could be temporally integrated to produce motion direction estimates significantly higher than chance levels (as estimated by permutation tests, **Figure 2B**). We observed substantially higher performance obtained from IT sites than V4 sites (%Δ (IT - V4) = 14.54%, paired t-test, p< 0.001; as estimated by integrating the neural signals from 0 to 600 ms post video onset) in predicting motion direction following the onset of the stimuli. Further analysis also revealed that the decoding accuracies obtained from IT sites scaled more rapidly with the number of sampled sites (**Figure 2C**; compare blue data points (IT) with magenta (V4)), suggesting a denser and possibly more precise representation of motion information in the IT cortex. Of note, these scaling analyses depend on several factors: video repetition number, the number of training examples per category, the total number of categories (e.g., motion directions), etc. Therefore, we strongly encourage the readers to consider only the relative comparisons between IT and V4 made here (performed under the exact training parameters) and not the absolute values. Next, we explored the effectiveness of an instantaneous (30 ms bins of neural responses) linear readout approach, deploying a linear classifier (linear discriminant analysis, LDA, classifier, *see Methods*) to decode motion direction from the same IT and V4 responses (across 192 sites from each area).

**Figure 2.**
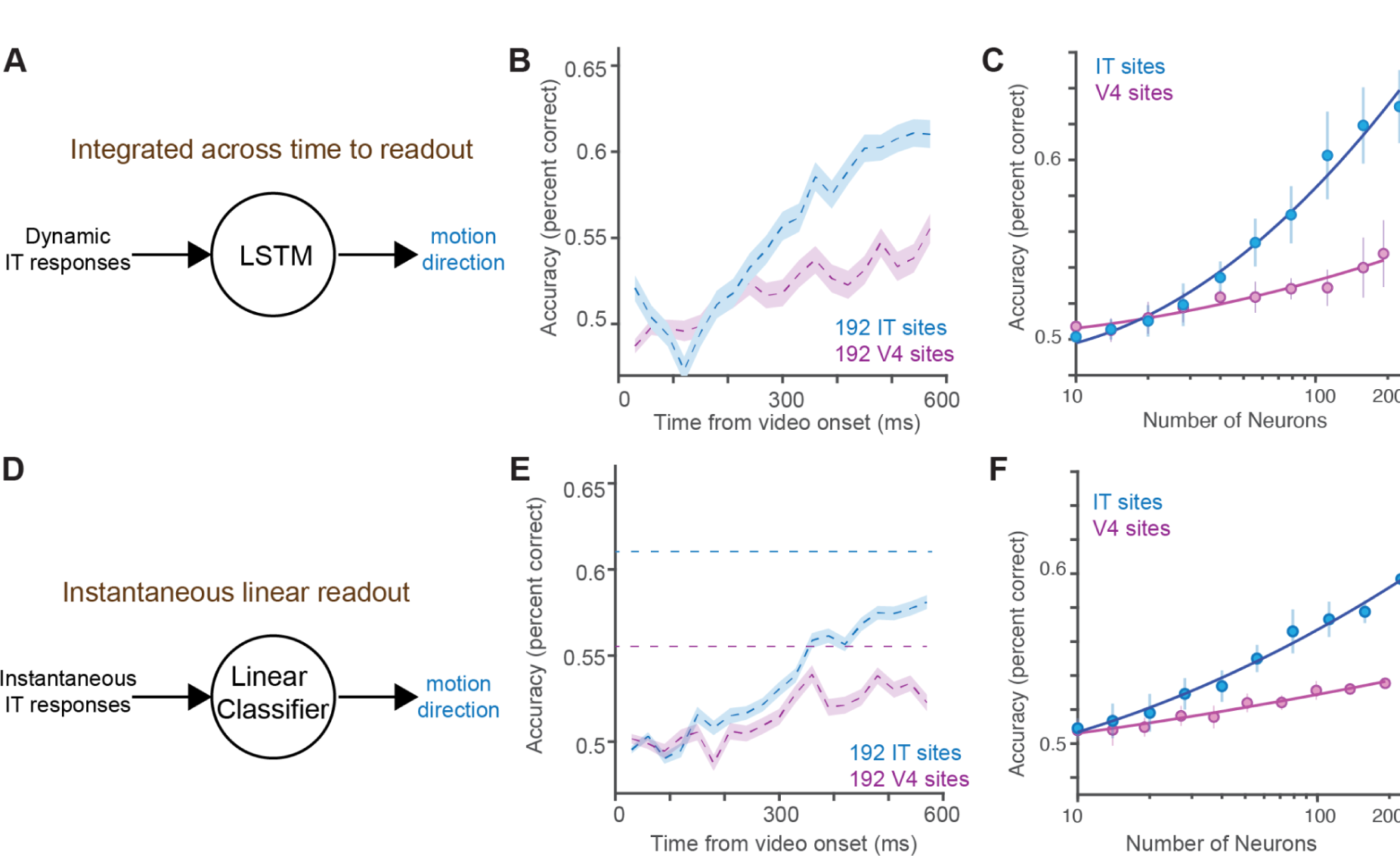
Comparative analysis of temporal and instantaneous decoding strategies in primate visual cortex for motion prediction. A. Schematic representation of a Long Short-Term Memory (LSTM) network processing dynamic IT responses to predict motion direction. This depicts the mechanism by which time-sequential neural data could be integrated (likely in an area downstream of the corresponding brain area) to form an accurate prediction of object movement. **B.** Accuracy of motion direction prediction over time using an LSTM decoder, comparing the performance of 192 IT (blue) and V4 (magenta) neural sites. The graph shows the temporal evolution of the decoding accuracy following the video onset, highlighting the differences between the two brain regions – IT decodes being significantly higher than V4 (%Δ (IT - V4) = 14.54%, paired-t-test, p< 0.001; as estimated by integrating the neural signals from 0 to 600 ms post video onset). **C.** Scaling of decoding accuracy with respect to the number of neurons from IT and V4 sites using the LSTM network. The plot demonstrates that increasing the number of neurons enhances the predictive capability, with IT sites showing a steeper accuracy gain. **D.** Schematic representation of an instantaneous linear readout strategy employing a linear classifier to decode motion direction from instantaneous IT responses. If accurate, this model would emphasize a direct, non-sequential approach to estimating motion direction – as a predictive signal contained within the IT population activity. **E.** Performance comparison of instantaneous linear readout for motion direction decoding between 192 IT (blue) and V4 (magenta) sites over time, with IT sites significantly outperforming V4 sites (%Δ (IT - V4) = 11.1%, paired t-test, p< 0.001; as estimated at the time window 550-600 ms post video onset). The blue and magenta dashed lines refer to the accuracies achieved by IT and V4 using LSTM-based readouts, respectively. **F.** Analysis of how decoding accuracy scales with the number of neurons in an instantaneous readout model. Similar to the LSTM approach, an increase in neuron sample size corresponds to better performance, particularly in IT sites.

### Object motion direction can be linearly decoded from V4 and IT cortex population activity

Motion signals (as direction or speed tuning functions) in the primary visual cortex ^33^ or areas like MT ^34^ can be estimated without nonlinearly integrating neural responses (as in **Figure 2A-C**) recorded from these neurons. If object motion velocity can be directly inferred with linear decoding algorithms – it would suggest that an upstream integration of spatiotemporal cues (like in areas V1 and MT) could be inherited in the V4 and IT cortices. We used linear discriminant analysis classifiers on the ventral stream activity to test how accurately we could read out the motion directions from these neural responses. Comparing the performance of linear decodes from 192 IT and V4 sites independently, we observed that both IT and V4 produced significant decoding accuracies (above 0.5, i.e., chance level), suggesting the presence of an instantaneous motion signal within the population responses in these areas. Similar to the LSTM results (**Figure 2B**), we again observed that the IT sites displayed significantly higher (%Δ (IT - V4) = 11.1%, paired-t-test, p< 0.001; as estimated at the time window 550-600 ms post video onset) decoding accuracies than V4 (**Figure 2E**), further affirming the robustness of the IT cortex’s predictive signal for motion direction. Interestingly, the non-linear integrative readouts (**Figure 2A-C**) produced a higher decoding accuracy (as indicated by the respective dashed lines in **Figure 2E**) than the linear readouts. This suggests that additional transformations of IT responses (likely in downstream regions) produce more linearly separable solutions for motion velocity readouts – a hypothesis to test in future studies. We also estimated how decoding accuracies scaled with the number of neurons within the instantaneous readout framework (**Figure 2F**). Consistent with the LSTM findings, increasing the neuronal sample size improved decoding performance (at a higher rate for IT than V4), underscoring the IT cortex as a pivotal region for interpreting visual motion cues. These observations collectively suggest that compared to area V4, the primate IT cortex harbors a more potent and scalable mechanism for predicting motion direction, whether through the integration of temporal sequences or instantaneous readouts. Hence, from hereon, we have primarily focused on IT-based decoding results.

### Object identity decodes precede object motion estimates

Most previous studies ^6,7,35^ almost exclusively test object identification capabilities from the IT cortex while presenting static images to the animals. To shed light on how the IT-based identity estimates of the objects tolerate the position shifts introduced during motion within the central 8 degrees of field of view, we further estimated the object identification accuracies from IT population activity using the non-linear (LSTM) and linear (LDA) decoding algorithms. We first observed that object identity information is accurately decoded using LSTM-based temporally integrative algorithms (**Figure 3A**), with significantly higher accuracies than the motion direction estimation (%Δ (identity - motion) = 50.3%, paired t-test, p< 0.001; as estimated at the time window 570-600 ms post video onset), using the same algorithms (compare red and blue curves in **Figure 3B**). Interestingly, the identity information was available much earlier than the direction estimates. Similar to object motion direction estimates, the identity estimates also scaled with neural sampling size – but reached higher accuracies with much fewer neurons (**Figure 3C**; red dashed and blue dashed lines showed the accuracies of the monkeys during an object identification and direction recognition task, respectively). The accuracy and neural sampling-based scaling of object identification were much weaker when estimated by linear classifiers **(Figure 3D, F**) compared to the LSTM-based decoders (**Figure 3E**, also compare red curves in **Figure 3B and E**, and **Figure 3C and F**). Given that object identity information was available at significantly lower latencies from the IT cortical responses, we next probed the criticality of this identity information for the motion direction estimates extractable from the IT population activity.

**Figure 3.**
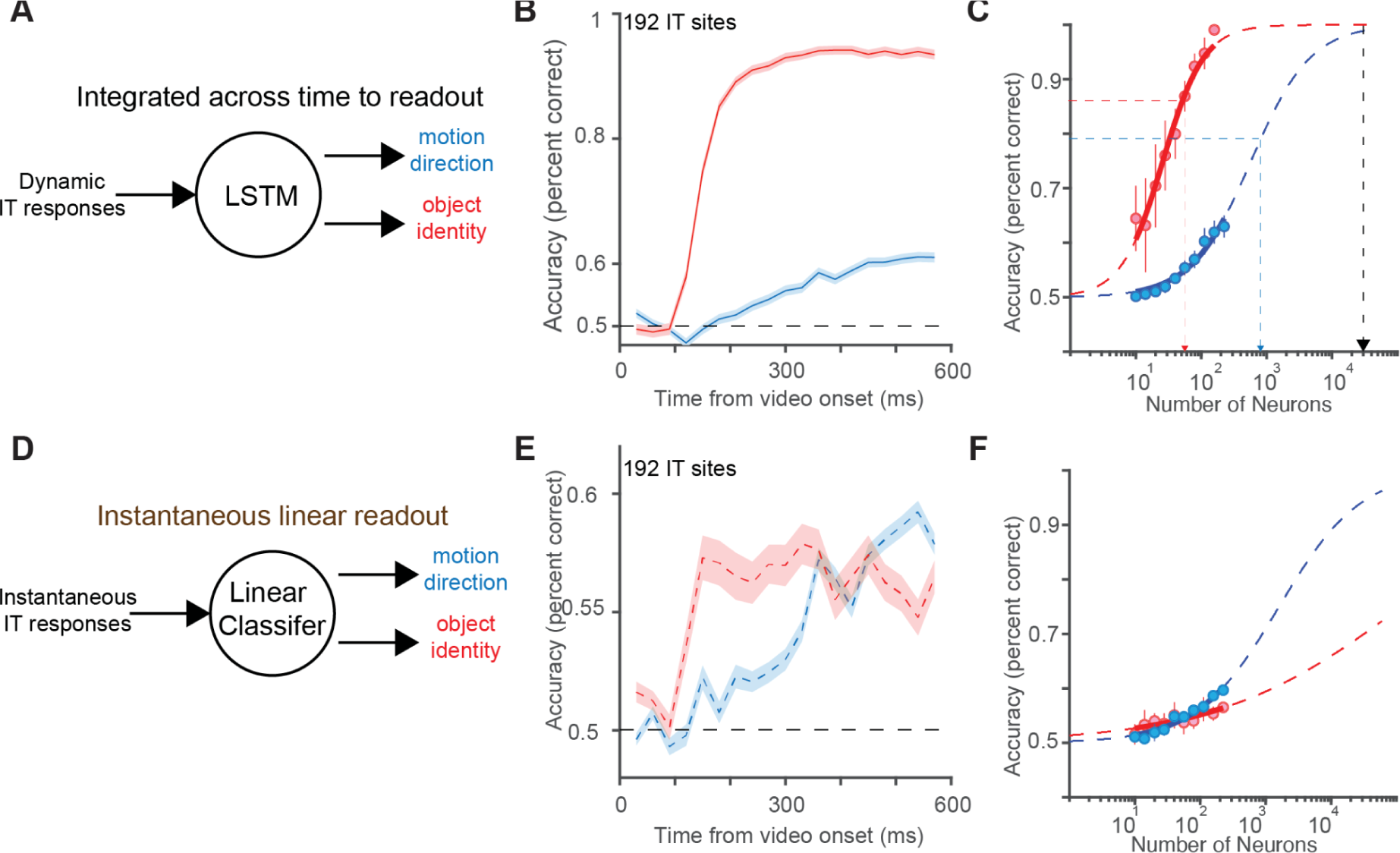
Comparing the readout of object identity and motion direction from the population activity in the IT cortex using temporal and instantaneous strategies. A. Schematic representation of a Long Short-Term Memory (LSTM) network processing dynamic IT responses to predict object identity and motion direction. **B.** The temporal decoding accuracy for object identity and motion direction from 192 IT sites as a function of time after video onset. The red curve indicates object identity, showing a faster and significantly higher accuracy (%Δ (identity - motion) = 50.3%, paired-t-test, p< 0.001; as estimated at the time window 570-600 ms post video onset) than the blue curve representing motion direction. **C.** Scaling analysis of decoding accuracy for object identity (red) and motion direction (blue) with the increasing number of IT neurons. The red curve achieves a higher accuracy more rapidly than the blue curve, indicating quicker access to object identity information in the IT cortex. **D.** Schematic representation of an instantaneous linear readout strategy employing a linear classifier to decode object identity and motion direction from instantaneous IT responses. **E.** Comparative decoding accuracy of object identity and motion direction using an instantaneous linear model with 192 IT sites over time. The red and blue curves show that object identity (red) is decoded with greater accuracy than motion direction shortly (∼120-150 ms) after the video begins, compared to motion direction (peaking > 300 ms). **F.** Scaling of decoding accuracy for object identity (in red; dashed line represents the extrapolation) and motion direction (in blue; dashed line represents the extrapolation) in a linear readout model. Unlike the LSTM-based decoders, we observe that object identity scales with a significantly shallower slope compared to motion direction across the number of neurons.

### Motion direction information within the IT cortex is tolerant to motion-preserving object appearance removal

Similar to most object tracking algorithms ^36^, where a segmented form of the identified object is often annotated and provided as a reference to be tracked, we observed that object identity estimates from the IT cortex precede (achieve higher accuracies under the same neural sampling regime and training set) the object motion direction estimates. Therefore, we next asked whether this information about the object’s appearance is necessary for the motion direction estimates from the IT cortex (**Figure 2-3**). To test that possibility, we removed the object appearance from 100 original videos (OV) without losing its motion content using a previously established technique^37^ (*see Methods*). We refer to the resulting videos as appearance-free videos (AFV). We trained two monkeys to perform a binary object discrimination task (*see Methods*, **Figure S1A**) and an 8-way direction estimation task *(see Methods*, **Figure S1B**) and also recorded IT activity (in 2 separate monkeys, 84 sites in total) while they passively viewed these two types of videos. First, our results reveal that when monkeys were tasked with identifying a target object that had been stripped of its appearance (AFV), their performance was significantly impaired and indistinguishable from chance level (**Figure 4B**, left panel) compared to their ability to identify the same object in the original videos (OV) containing appearance information. However, this decrement in performance was not observed when the task required discriminating the direction of motion: The monkeys’ performance remained high and statistically indistinguishable whether the object’s appearance was present (OV) or absent (AFV), suggesting that the perception of motion direction can be behaviorally dissociated from the object’s appearance. Next, we asked whether the IT-population-based decodes of object identity (as shown in **Figure 3A**; LSTM-based decoders) are consistent with the behavioral patterns. Indeed, we observed that removing the object appearance significantly reduced the neural-based decoding accuracy to chance level for the AFV compared to those for the OV (**Figure 4D**). Interestingly, we could decode the motion direction in the video significantly above chance for both OV and AFV. We observed significant accuracies for both LSTM-based and linear-classification-based approaches. Notably, the decoders trained (i.e., the same set of weights and biases) with AFV data also generalized to OV test set responses and vice versa (**Figure S2**), demonstrating that the same motion signal processing in the IT cortex can operate without the presence of an object.

**Figure 4.**
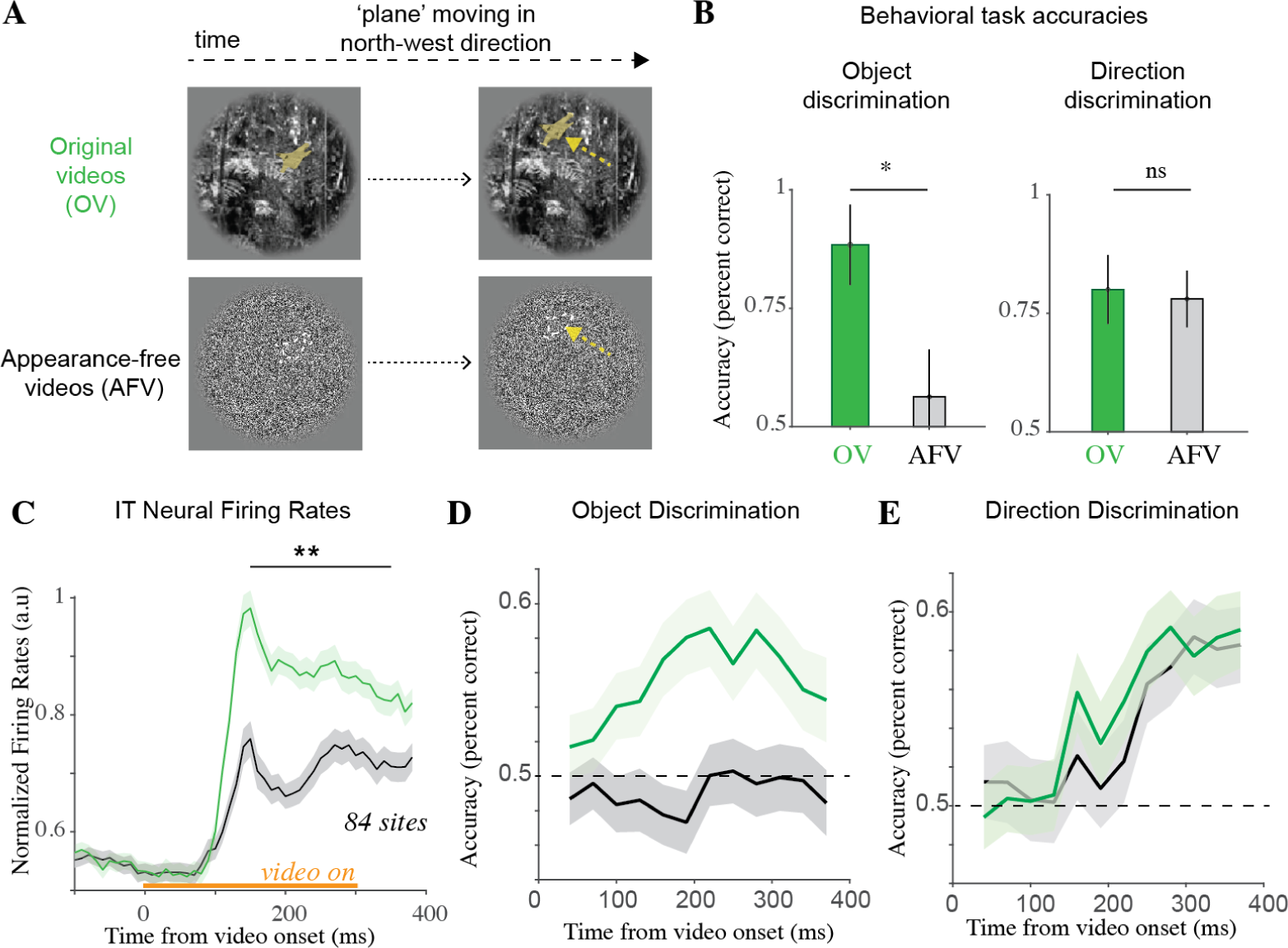
Decoding motion directions from IT population activity for videos with and without object appearance. A. The top panel shows two frames from a video (300 ms) of a plane moving in the northwest direction – the original videos (OV). The yellow highlighting of the plane is not part of the original video (shown here only for illustration). The bottom panel shows the corresponding appearance-free version of these video frames, produced by methods described in Ilic et al. ^37^ – referred to as the appearance-free videos (AFV). The white outline of the plane is not part of the video; we outlined it here for comparison to the above video. **B.** Monkeys (n=2) behavioral accuracies on two tasks performed on these videos. During the object discrimination task (left panel), the video containing a target object (with or without the appearance) is followed by a choice screen with two options (one target object, another distractor object). The monkeys were required to choose the target object shown in the video (see Supplementary Material). As shown here, monkeys performed significantly better during the OV than their appearance-free counterparts (Δ OV-AFV = 0.29, paired t-test, p<0.0001), which were indistinguishable from chance-level performance. During the direction discrimination task (right panel), monkeys were trained to saccade to one of the eight fixed locations (see **Figure S1**) based on the perceived direction of motion of the objects. We observed that monkeys showed equally high performance (significantly indistinguishable) for both OV and AFV. **C.** IT neural firing rates (33 sites in monkey 1 and 51 sites in monkey 2) time averaged across AFV and OV. As expected, appearance-based videos (OV) drove the IT population’s evoked responses significantly higher (%ΔIT response (OV - AFV) = 35.05%, t(83) = 8.32, p<0.001, computed at latency 150-200 ms post video onset) than appearance-free videos (AFV). **D.** Decoding of object identity from the monkey IT population activity. LSTM-based (trained and tested in a cross-validated way on OV and AFV responses independently) decoding models linking IT population activity to object identity (similar to Figure 3B) discrimination perform well above chance levels (p<0.05, permutation test, 270-300 ms post video onset) for OV but are indistinguishable from chance-level for AFV. **E.** Decoding of motion direction from the monkey IT population activity. LSTM-based (trained and tested in a cross-validated way on OV and AFV) decoding models linking IT population activity to direction discrimination perform well above chance levels (p<0.05, permutation test, 270-300 ms post video onset) >200 ms post video onset for both AFV and OV. Errorbar bands denote s.e.m across videos.

### IT population activity can predict object speed

Object motion also comprises another important variable in addition to the direction of the moving object – which is its speed. Therefore, we next asked whether an object’s speed can also be decoded from the IT cortical responses using non-linear, temporally integrative (LSTM-based, **Figure 5A**), and linear regression-based (**Figure 5B**) methods. We recorded large-scale activity across the V4 and IT cortices in two macaques while they passively fixated on brief (300 ms) videos presented at the central 8 deg. Similar to the direction estimates, we observed that we could decode object speeds with significant accuracies (greater than chance-level as estimated by permutation tests, *see Methods*) from the distributed population activity of the IT cortex. Consistent with earlier results on motion direction estimates, we observed that the decoding based on V4 population activity was, in general, weaker than the ones derived from the IT cortex (%Δ (IT - V4) = 16.8%, paired t-test, p< 0.001; as estimated at the time window 270-300 ms post video onset). The ventral visual pathway is often associated with the processing of slower foveal information, whereas the dorsal visual pathway is associated more with fast peripheral information ^38,39^. Indeed, consistent with this functional distinction, we observed that IT responses to slower speeds (e.g., <10 deg/s) were higher than with higher speeds (**Figure 5C**; top inset). In addition, we could also better decode (with more accuracy) lower speeds than higher speeds (**Figure 5C)**, and the accuracies were significantly negatively correlated with object speed (Pearson R = -0.39, p = 0.01).

**Figure 5.**
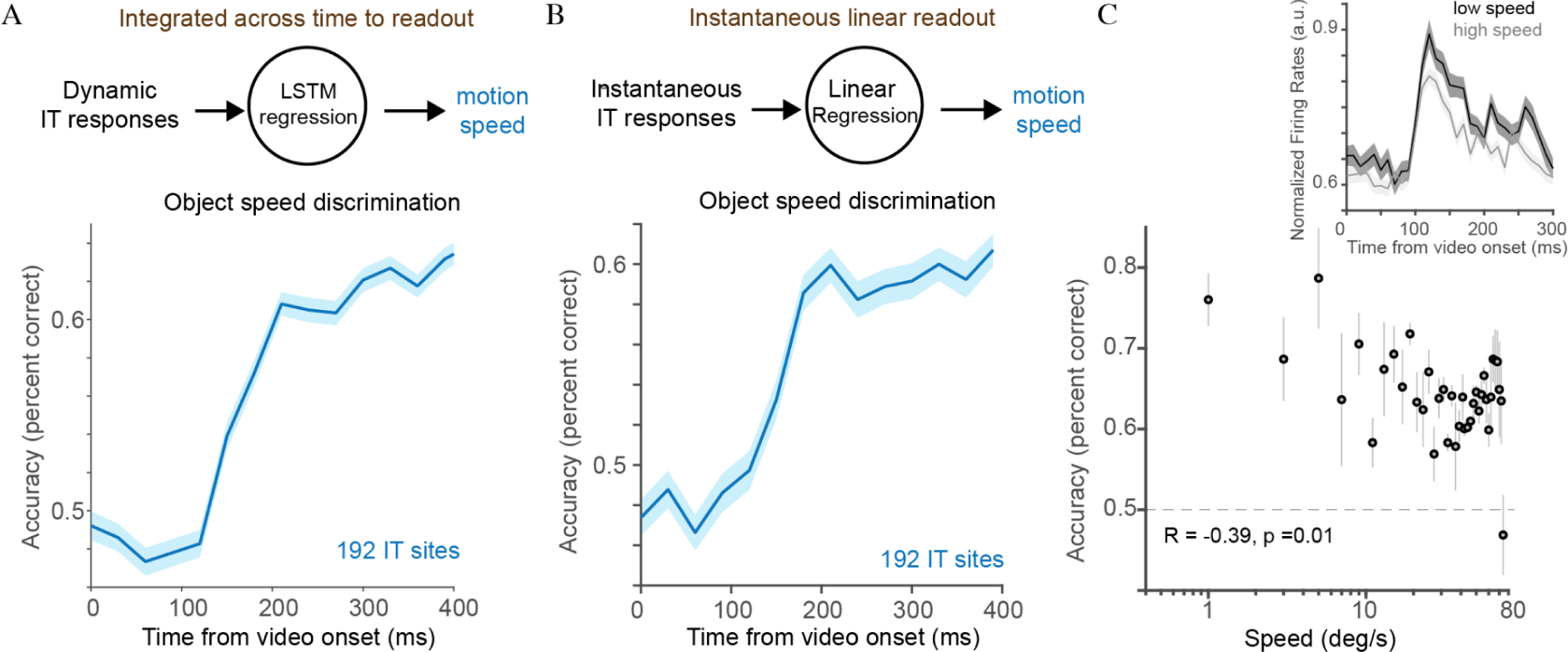
Decoding object motion speed from IT population activity. A. The accuracy of speed discrimination over time using an LSTM network that receives the dynamic IT responses as input, with a sample size of 192 IT sites. The bottom panel shows that the resulting decoding accuracies (cross-validated across videos) are significantly greater than the chance level (p<0.05, permutation test, at time bins > 150 ms). The shaded area represents the standard error of the mean across videos. **B.** The accuracy of an instantaneous linear classifier using independent 30 ms bins of IT responses from 0 ms post video onset (using 192 IT sites), with the shaded area indicating the standard error of the mean across videos. Similar to the LSTM readout, decoding accuracy reaches a level significantly higher than chance, approximately 150 ms after video onset. **C.** The relationship between object speed (degrees/s) and decoding accuracy. We could better decode (with more accuracy) lower speeds than higher speeds, and the accuracies were significantly negatively correlated with object speed (Pearson R = -0.39, p = 0.01). Inset: Neural response time course for low versus high-speed stimuli, highlighting the differential activation over time (higher average firing rates for lower speeds).

### IT population activity-based decodes predict object identity-dependent speed discrimination behavior

So far, our results have provided evidence for decodable object motion information within the ventral stream population activity, specifically within the IT cortex of the macaques during passive viewing of the videos. However, is this signal just an epiphenomena, or can this be linked to the pattern of responses measured during object-motion-dependent behaviors? To test the behavioral linkage, we first designed an object-dependent speed discrimination task. As shown in **Figure 6A**, monkeys (n=2) fixated on a cross for 100 ms to initiate a trial, after which we presented a 300 ms video containing two objects (one moving faster than the other). After a blank period of 100 ms, we presented canonical images of those two objects, and the monkeys had to select the one that moved faster. In this task, the monkeys achieved high accuracy (average accuracy = 0.82 ± 13, Mean ± standard deviation, percent correct, **Figure 6B**) and showed reliable behavior (trial split-half reliability of estimating accuracy per image = 0.89). We also recorded the neural responses to these videos across the IT cortex during passive viewing of the 100 Test videos in two task-naive macaques. Cross-validated linear decoding of the task accuracies (*see Methods*) shows that neural responses averaged between 340-410 ms post-video onset produced decoding patterns that were significantly consistent (noise corrected R = 0.6, 192 IT sites, estimated at 340-410 ms post-video onset, **Figure 6C**) with the behavioral patterns. As demonstrated earlier, these estimates are limited by the neural sampling size of the experiments and show a general trend of improvement with the number of neurons (as estimated by varying the number of neural sites sampled randomly from 2 to 120, **Figure 6D**) – establishing the possibility of a highly critical functional role of the IT cortex in these behaviors.

**Figure 6.**
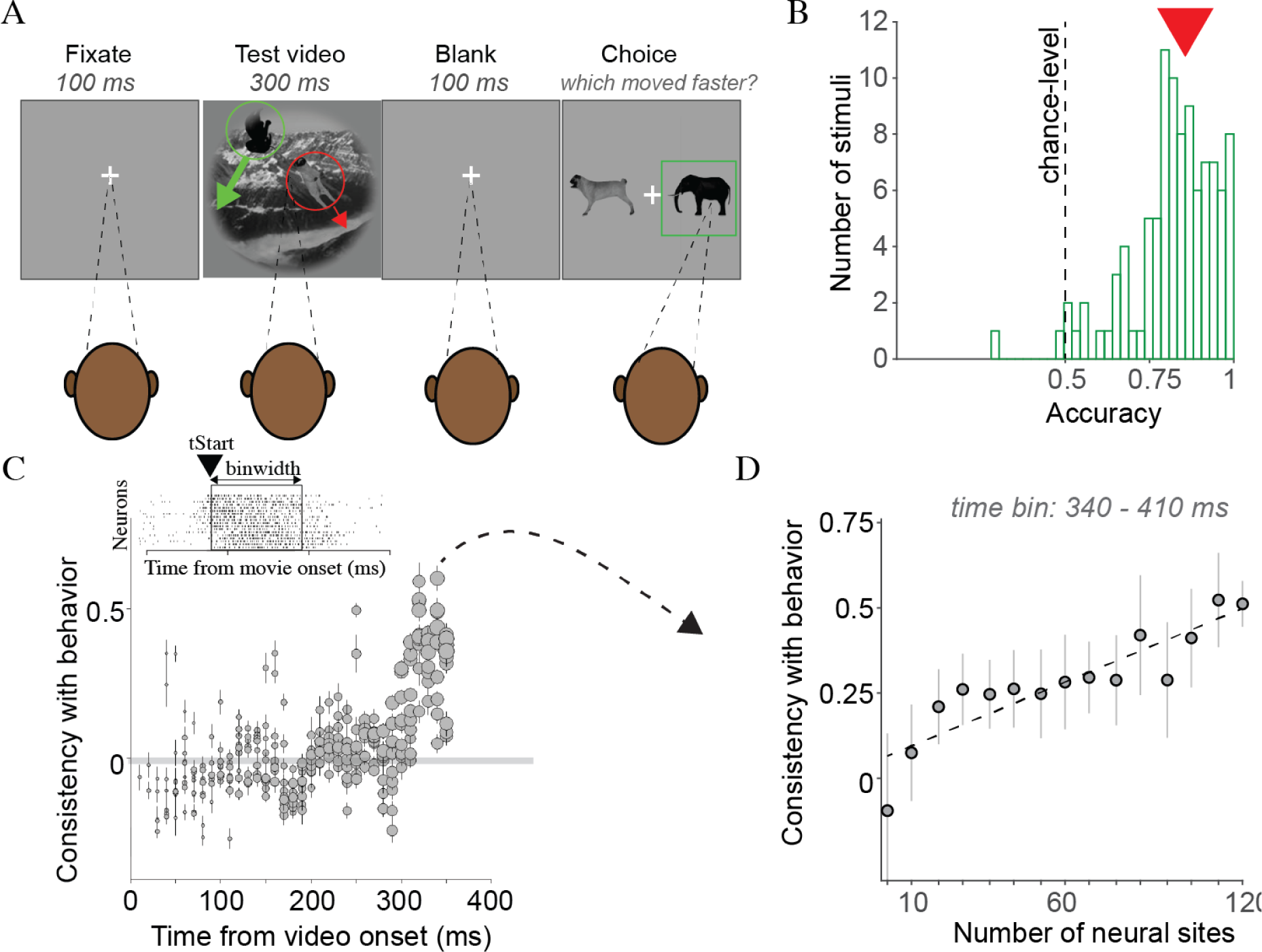
Linking IT population activity with speed discrimination behavior. A. Speed discrimination task. Monkeys (n=2) fixated on a cross for 100 ms to initiate a trial, after which we presented a 300 ms video containing two objects (one moving faster than the other). After a blank period of 100 ms, we presented canonical images of those two objects, and the monkeys had to select the one that moved faster. **B.** Distribution of accuracies of the monkeys’ behavior (averaged per video) across 100 videos. The dashed line indicates the chance level of accuracy, and the red arrow shows the average accuracy (0.82 ± 13, mean ± standard deviation). **C.** Consistency of IT-based neural decodes of the velocity discrimination task (*see Methods*) across multiple choices of temporal decoder parameters (these were selected from two free parameters: time start and bandwidth with respect to the onset of the video, as schematically demonstrated in the inset). The highest consistencies were observed for signals integrated between 340 to 410 ms post video onset (noise corrected R = 0.6, 192 IT sites). **D.** Analysis of the relationship between the number of neural sites sampled and the consistency of their collective decoding with the monkeys’ behavioral responses. The selected time bin (340-410 ms post-video onset) that yielded the highest consistency is highlighted, with the trend line indicating improved consistency with an increased number of neural sites.

### Microstimulation of IT provides causal evidence for the critical role of the IT cortex in the speed discrimination task

While the correlation of behavioral patterns and neural decoding accuracies strengthens the possibility of a functional role of the IT cortex in these behaviors, it is not a direct causal test of this linkage. Therefore, we performed microstimulation of the IT cortex (to disrupt its activity as demonstrated in previous studies^40,41^ temporarily) in two macaques (left and right hemispheres in each monkey, respectively) during the tasks to test whether the animal produces specific contralateral deficits implicating the causal role of IT. We applied 10 uA of bipolar pulses for 150 ms (starting 50 ms post-video onset) across 16 randomly chosen sites on the anterior array. The stimulation pulses were biphasic, with the cathodal pulse leading. Each pulse was 0.2 ms in duration, with 0.1 ms between the cathodal and anodal phases. On each trial, the objects started from two distinct positions on opposite hemifields, and the movement of the two objects was primarily present in their respective hemifields (with minor cross-overs). We specifically implemented this aspect in the design to test the contralateral nature of the microstimulation. Microstimulation significantly reduced accuracy in the speed discrimination task for both ipsilateral (p<0.001, t(99)=9.53, paired t-test) and contralateral targets (p<0.001, t(99)=5.33, paired t-test). However, importantly, consistent with the known lateralization of the IT cortex ^42^, we observed a more pronounced effect (Δ for ipsi and contra shown in the inset) on the accuracy of detecting contralateral versus ipsilateral targets (p<0.001, t(99)= 4.52, paired t-test).

### IT population activity predicts object-motion-based future event prediction behavior

In the speed discrimination task described above, the behavioral report can be accurately estimated based on the information presented during the video. It does not require a prediction of the future based on those estimations. We designed a future event detection task to ask whether IT-based motion estimates could be used to make forward predictions of events. This task links our study to ideas about an intuitive physics engine running in the brain ^43^, as often proposed in the cognitive sciences. We presented the monkeys with a 300 ms video containing a ball and a car moving at different speeds. The ball moved vertically downwards, and the car moved horizontally to the right (**Figure 8A**). Depending on their velocities, they would either collide (*condition 1*), the ball would move past the car (*condition 2*), or the car would move past the ball (*condition 3*). Following a 100 ms blank screen after the Test video, the monkeys were shown two images on the choice screen depicting two possible conditions (one likely and one unlikely future event). Given the brief video, their task was to choose the likely outcome of the video from the choice frames within the next 1500 ms. The videos were always presented such that they were shorter than the full event. We tested the animals (n=2) on three phases of the task. In phase 1, we showed the animals a training set of 400 videos (full version; 700 ms duration per video) while they passively fixated on these videos. The primary purpose was to make them aware of the possibilities in these videos (i.e., familiarization with conditions 1-3). Next, on this subset of 400 videos (training set), we trained the animals to perform the future event detection task. Lastly, we used a held-out video (test) set with 105 videos to measure their behavioral reports. On average, the monkeys showed an accuracy of 0.68 (significantly higher than chance-level; t-test, t(104) = 13.5, p<0.001; **Figure 8B**). On the other hand, we recorded spiking activity across the IT cortex (192 IT sites) in 2 task-naive macaques while they passively viewed the shortened (300 ms) Test videos. We observed that consistent with previous results, the IT-based linear decodes could attain monkey-like significant high accuracies in these tasks, which peaked around 200 ms post video onset (**Figure 8C**). Similar to the analyses for **Figure 6C**, we performed a cross-validated search to identify the most behaviorally aligned decoder parameter. Interestingly, the IT-based predictions from a temporal integration window of 350-400 ms post video onset across videos showed the highest significant correlation (Pearson R=0.4, p<0.001, permutation test) with the measured behavioral accuracies. This time window is very similar to the time window that produces the best match for the speed discrimination task (**Figure 6C**) – suggesting that the macaque brain might rely on the temporal code most relevant for speed estimates to make future predictions of objects’ trajectories. **Figure 8D** demonstrates how the correlations for this 50 ms time bin evolve (the gray-shaded region refers to the null distribution computed by shuffling the condition labels, re-estimating the decoding accuracies, and correlating them with the behavioral data).

## Discussion

In this study, we highlighted the critical role of the ventral stream circuits in primate object motion perception. Our findings revealed that neural signals related to object motion (direction and speed) are present in the distributed population activity across the macaque ventral stream — the ones in the IT cortex being more easily decodable with fewer neurons than those in area V4. We also demonstrated that these signals accurately predict the behavioral patterns measurable in monkeys during object motion speed discrimination tasks. Moreover, by demonstrating behavioral deficits induced by perturbation of the IT cortex with microstimulation, we further confirmed the critical functional role of the IT responses in generating accurate behavioral reports during motion-related tasks. In the following sections, we will discuss our results and their broader implications and limitations.

### The ventral stream houses key circuits relevant to object motion perception

Our main finding underscores the ventral stream, specifically areas V4, and IT, as critical to object motion perception in macaques. This insight aligns with the established role of the ventral stream in visual object recognition and suggests its involvement extends further to the domain of motion perception. However, the degree to which foveal object motion processing is exclusive to the ventral stream remains an open question. While our results indicate that the IT decodes slower object speeds more accurately, suggesting a ventral stream specialization, two observations invite further scrutiny into possible dorsal stream contributions. First, we observed that decoding of fast object speeds (typically attributed to the dorsal stream) also remains robust, surpassing chance levels (**Figure 5C**). Second, motion direction information from appearance-free videos could be extracted with the same neural population code as object-based videos (**Figure 4**). These observations hint at a more complex interplay between ventral and dorsal visual pathways in motion perception than previously understood. Given our experimental results, a series of causal experiments can now be performed to disentangle the contributions of these two streams. For instance, manipulating activity in dorsal areas like MT, MST, and LIP while recording the response of V4 and IT cortex to the same videos as used in this study could reveal the functional role of the dorsal regions in shaping the object motion representation in the ventral stream. Additionally, computational models with varying architectures can be instantiated to offer insights into the potential circuits and computations that lead to these ventral stream representations. By simulating different aspects of both streams, we can better hypothesize how the brain might integrate information from these two pathways to perceive objects in motion.

While our study provides a systematic characterization of object motion decodability from the ventral pathways, there are several prior reports of motion-related responses in the ventral stream ^28–31,44,45^, especially when using stimuli much more complex than those typically used in the studies of the dorsal stream (such as random dot motion). Does the ventral stream mechanism process motion de novo (or independently)or augment the preprocessed motion information available in the dorsal neurons? There might be three distinct sources providing the initial motion-related signals to the ventral stream for integrating it with object identity information:

1. input from dorsal stream areas such as MT via projections into area V4: Previous studies have shown that not only neurons in the visual area V4 can be responsive to kinetic gratings ^46^, but they may also form a motion direction preference map ^47^, most likely inherited through monosynaptic connections from area MT ^48,49^. Indeed, the hierarchical organization of the ventral stream ^50^ would allow such direction-selective signals in V4 to feed into the IT cortex.
2. a subcortical input: There also is evidence for a subcortical motion pathway with the pulvinar playing a major role ^44^. The pulvinar, which is heavily connected to MT and MST ^26,51,52^, contains a significant number of direction-selective neurons ^53^ and projects to ventral stream areas, such as V4, TE, and TEO ^26,54,55^. Therefore, the ventral stream motion responses can be presumed to reflect an injection of pulvinar motion information.
3. Feedback originated at higher-order areas such as the prefrontal cortex: Higher-level cognitive processes like attention, memory, and prediction may influence the IT cortex’s response to motion. Direct connections exist between the prefrontal cortex (PFC) and the ventral stream sites we studied ^56–58^. Notably, the PFC shows a substantial presence of direction-selective responses ^59^, similar to neurons in area MT. These results suggest that ventral stream motion-related signals might originate at the PFC level, facilitating the selection of objects based on their movement direction. This hypothesis contrasts with a pure sensory analysis of basic visual motion in the dorsal stream.

As mentioned above, a combination of model development with varied hypothesized architectures ^12,13^ and goals^60^ along with targeted brain perturbation studies^61^ are required to fully discriminate among these alternatives (while a combination of all three cases is also plausible).

### Causal evidence for the role of IT in object motion perception

The object motion-related signals we observed in the ventral stream are not epiphenomenal, as demonstrated by the causal disruption of information processing in the IT cortex via microstimulation, which impairs the monkeys’ performance in a speed discrimination task (**Figure 7**). Given that object processing in IT can be disrupted by electrical, pharmacological, or optogenetic stimulation ^23,62^, one explanation of our results can be that microstimulation of the IT cortex produces an object-specific general deficit across all task conditions (unrelated to its motion), and this results in the measured behavioral performance changes. However, if this were the case, we would also expect to find a speed-independent disruption in behavior when the contralateral hemisphere is stimulated. Our results show a significantly higher deficit when the faster-moving (Target) object is in the contralateral hemifield – supporting the hypothesis that stimulation impaired the combined signal of object motion speed and object identity (aka the “binding”). Our findings, which suggest that ventral stream motion signals play a crucial role in shaping behavior when selecting relevant objects, are consistent with results observed in other studies demonstrating that lesioning ventral stream areas like V4 or TEO reduces monkeys’ ability to filter out distractor information across various feature domains, including motion ^63,64^.

**Figure 7.**
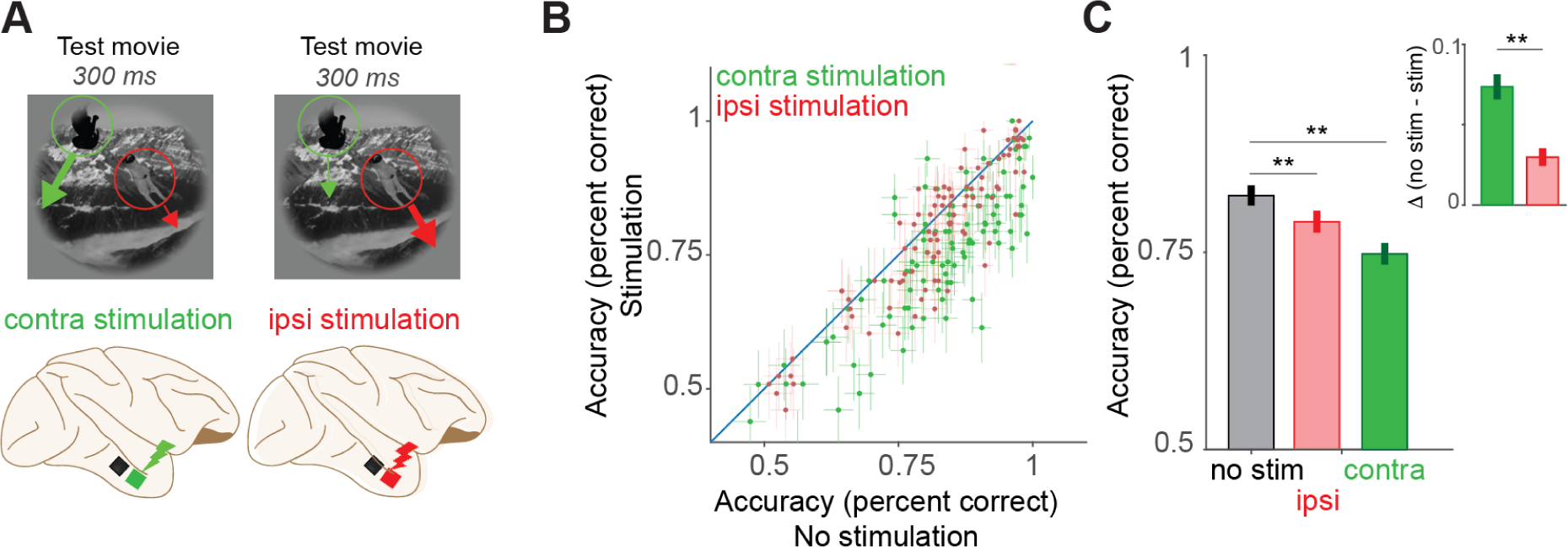
Causal test of the role of IT in object motion speed discrimination. A. Microstimulation Conditions. We stimulated in two primary conditions – ipsilateral (when the target object that moved faster was in the ipsilateral hemifield compared to the site of stimulation) and contralateral (when the target object that moved faster was in the contralateral hemifield compared to the site of stimulation). Microstimulation (10 µA) was always applied across 16 randomly chosen sites on the anterior array. **B.** Comparison of behavioral accuracy video-by-video (each dot refers to a video) on the task (see Figure 6A) during no stimulation trials and stimulation trials for the contra (green) and ipsi (red) conditions. **C.** Microstimulation significantly reduced accuracy in the speed discrimination task for both ipsilateral (p<0.001, t(99)=9.53, paired t-test) and contralateral targets (p<0.001, t(99)=5.33, paired t-test). We observed a more pronounced effect (Δ for ipsi and contra shown in the inset) on the accuracy of detecting contralateral versus ipsilateral targets (p<0.001, t(99)= 4.52, paired t-test).

**Figure 8.**
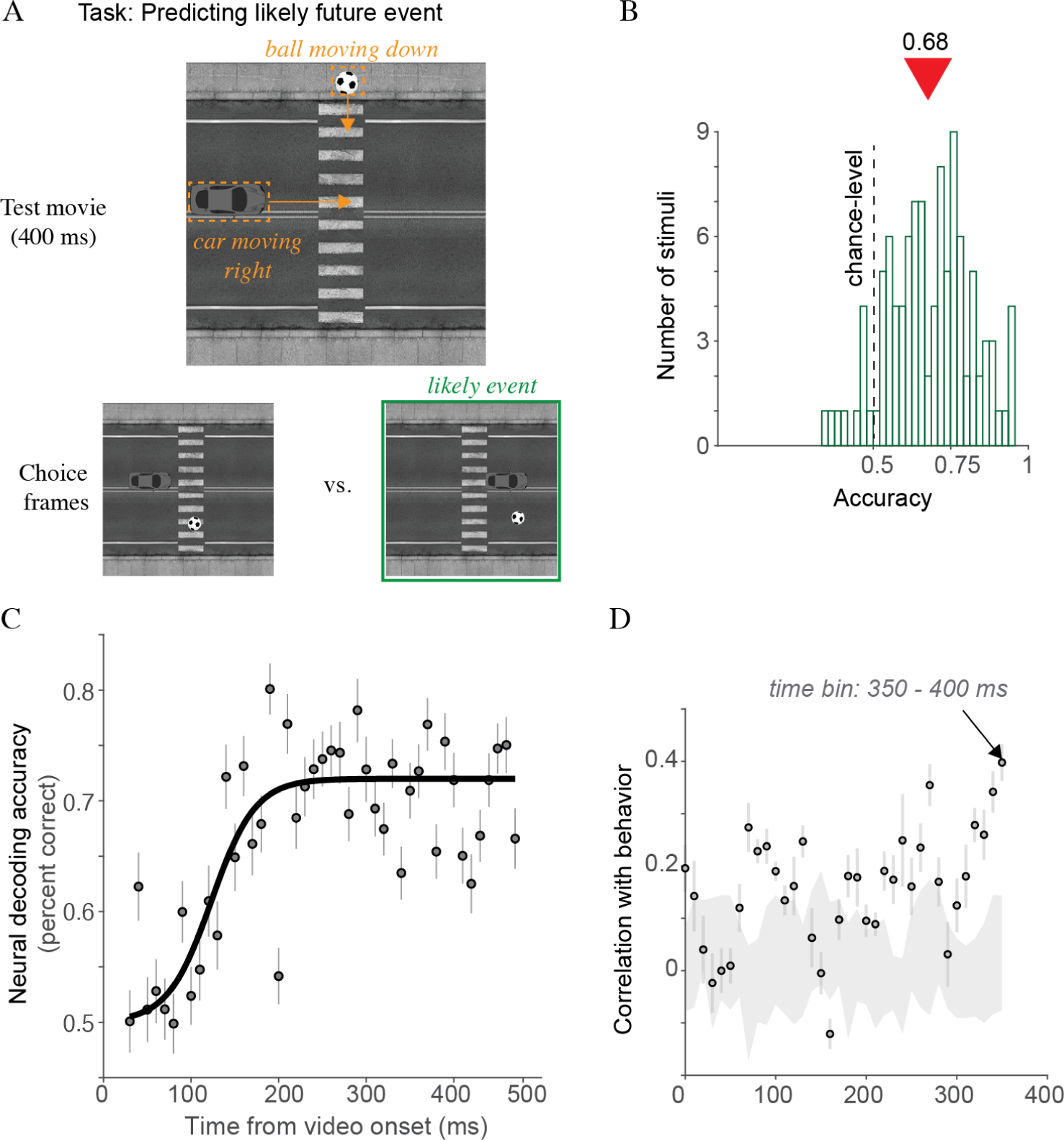
Linking IT population activity with speed-based future event prediction task. A. Future event prediction task. Monkeys viewed a 300 ms video containing a moving ball and a car. Both objects moved at varying speeds. The ball moved vertically downwards, and the car moved horizontally to the right. Depending on their velocities, they would either collide (condition 1), the ball would move past the car (condition 2), or the car would move past the ball (condition 3). The videos were always shorter than the full event. The monkeys were shown two images on the choice screen depicting two possible conditions. Given the brief video, they had to choose the likely outcome in the video. **B.** Distribution of accuracies of the monkeys’ behavior (averaged per video) across 105 videos. The dashed line indicates the chance level of accuracy, and the red arrow shows the average accuracy (0.68 ± 13, mean ± standard deviation). **C.** The ability of population activity of IT was measured during the video to make predictions about the outcome of the task. The IT-based linear decodes could attain monkey-like significant high accuracies in these tasks, which peaked around 200 ms post video onset **D.** The correlation of the IT-decoder (timebin of 50 ms) output accuracies per video with those measured from the monkeys. The decodes estimated from the population activity measured between 350-400 ms post-video onset showed the highest consistency (Spearman R=0.4) with the monkey’s behavior. Errorbars denote standard deviation across cross-validation splits. The gray-shaded region refers to the null distribution computed by shuffling the condition labels, re-estimating the decoding accuracies, and correlating them with the behavioral data.

### Interpreting the timing differences in estimating object identity and motion

In our study, we consistently observed that object identity, whether read out instantaneously or through temporal integration, was estimated more rapidly than the velocity of object motion (see **Figure 3, Figure S3**). This latency differential may arise from several factors. One hypothesis posits that early identification of object identity is imperative for formulating predictions about object motion, necessitating that identity estimates precede motion velocity estimation. However, our analyses contrasting AFV and OV decoding (see **Figure 4**) do not substantiate this notion. Another conceivable explanation involves the processing of motion in the V4 and IT cortices, which could be facilitated by top-down feedback from higher cortical areas, such as the prefrontal cortex (PFC), or indirectly from dorsal stream regions. Our current dataset does not provide definitive evidence to refute this hypothesis. To elucidate the potential influence of these feedback mechanisms, future research should incorporate targeted recordings from prefrontal regions and employ causal methods to interrogate these networks. A further explanation we consider is that the observed results may reflect the inherent dimensionality of the tasks—object identification might intrinsically be a lower-dimensional problem compared to velocity estimation. This possibility is suggested by our sample scaling analysis (**Figure 3E, F**), which indicates that a greater number of neurons may be required for accurate object motion velocity estimates, resulting in an inference of increased latency within our sample-limited analyses. To extend our understanding beyond the data collected from IT cortices, artificial neural network models that are designed to operate with two streams offer promising avenues for further investigation. In these models, one stream is trained for single frames or relatively low temporal processing rates, while a second stream is trained for optical flow or relatively high temporal processing rates (for example, see ^65,66^. These models could provide additional insights into the computational mechanisms underlying these processes and potentially enable strategies to discriminate amongst these hypotheses.

### Dynamic stimuli processing offers insights into integration mechanisms also relevant for object recognition

In this study, we made another critical observation: object identification using instantaneous readouts, while adequate for perceiving static images as previously suggested ^8^, was significantly less effective (based on lower observed decoding accuracies) than temporally integrative readouts that accumulate information from IT responses over time (refer to **Figure 3B, D**). This discrepancy underscores the possible role of spatiotemporal integration in the IT cortex, where signals are processed over a duration. Contrasting with earlier studies ^7,8^ where static images were displayed for 100 ms, the frames in our dynamic stimuli were presented for approximately 33 ms. This shorter time frame likely contributed to the diminished decoding performance we observed with linear classifiers. These findings imply that temporal integration is not just a byproduct but a fundamental aspect of the computations performed in the IT cortex, particularly under conditions that mimic the dynamic nature of natural vision. Given these findings, we propose that for object recognition tasks, LSTM-based decoders may offer a decoding strategy that better aligns with behavior (see ^13^), even when applied to static images. This is predicated on the idea that temporal integration is a continuous process in the visual cortex, essential for interpreting both static and dynamic scenes. To further test our speculation, future studies could explore using LSTM-based decoding in static image recognition tasks. Such research would potentially validate the goodness of temporal integration strategies and refine our understanding of how visual information is processed in the brain for decision-making, whether the stimuli are stationary or in motion.

### Future constraints and insights for the next generation of ventral stream models

The integration of our study’s findings into developing the next generation artificial neural network (ANN) models of the ventral stream promise a transformative shift in how dynamic visual information is processed in the computational models. The ability to decode motion parameters and future predictions directly from the IT cortex offers a unique biological blueprint for future computational models. This suggests that incorporating ventral stream-like features into ANNs is not only desirable but critical for achieving a more nuanced understanding of dynamic visual processing. Current and future ANNs might therefore, achieve a higher degree of biological fidelity by incorporating ventral stream-like features. For example, our results (**Figure 4**) suggest a crucial divergence of IT responses from how current action recognition models might behave. Previous studies ^37^ have shown that ANN performance during action recognition often deteriorates under the challenge of image perturbations that obscure object appearance. The capability of the IT cortex to faithfully represent motion parameters and make predictions based on appearance-free videos, without the reliance on object appearance-based features, provides a compelling argument for rethinking ANN design with inspiration from the ventral stream visual processing. By presenting neural data that elucidate how various read-out algorithms can help interpret object motion based on IT population activity, we offer a dual-purpose tool. First, this data can be converted into a rigorous benchmark for evaluating the prediction accuracy of a new class of dynamic ventral stream models (in benchmarking platforms such as Brain-Score ^67^). Second, our data can also be used directly to augment the training (see ^68^) of existing models that support video processing.

### Data and Code Availability

All of the data and code to replicate the results and support future research directions will be available on https://github.com/vital-kolab/object-motion upon publication of the article.

## Methods

### Subjects

#### Non-human primates

The nonhuman subjects in our experiments were six adult male rhesus monkeys (*Macaca mulatta*). All data were collected, and animal procedures were performed, in accordance with the NIH guidelines, the Massachusetts Institute of Technology Committee on Animal Care, and the guidelines of the Canadian Council on Animal Care on the use of laboratory animals and were also approved by the York University Animal Care Committee.

### Visual Stimuli

High-quality images of single objects were generated using free ray-tracing software (http://www.povray.org), similar to a previous study ^8,69^. Each image consisted of a two-dimensional (2D) projection of a three-dimensional (3D) model (purchased from Dosch Design and TurboSquid) added to a random background. The ten objects chosen were bear, elephant, face, apple, car, dog, chair, plane, bird, and zebra. By varying six viewing parameters, we explored three types of identity while preserving object variation: position (*x* and *y*), rotation (*x*, *y*, and *z*), and size. All image frames were achromatic with a native resolution of 256 × 256 pixels.

#### Generation of videos for IT-based object motion speed and direction readout estimation

We first rendered the 2D images of the objects and pasted them on uncorrelated backgrounds as mentioned above. Then, based on the chosen speed (degree/s derived from pixels/s given 8-degree 256 x 256 frames), and the direction, the object was moved an adequate number of pixels programmatically in MATLAB while keeping the background stationary.

#### Generation of videos for the object motion speed discrimination task

For the speed discrimination tasks, we first selected the 2D rendered versions of two objects (a *target object that will move faster and a distractor* object that will move slower). We then randomly selected two attributes for these objects. The first one was which object would be placed on the right hemifield and which would be placed on the left hemifield. Then, we randomly selected a motion direction for each of these objects. Based on the motion directions, we decided whether the object would be placed toward the periphery of the image or the center. If the motion trajectory places the object out of the frame (e.g., leftward motion for an object placed on the right edge of the image), we choose the opposite direction for that object. Once the initial position, the motion direction, and speed (also assigned randomly) were determined, we added the objects on top of an uncorrelated naturalistic background. We moved the objects at each frame with the pixel shifts that were appropriately adjusted for their directions and velocity choices while keeping the background stationary. Using this approach, we generated 100 videos, 10 per target object.

#### Generation of videos for the future event prediction task

These videos were generated in MATLAB using three main images – 1) a ball, 2) a car, and 3) a background image of a road crossing. For each video, the background image remained stationary. The position of the ball was initialized at the top (of the y-axis) and center (of the x-axis), and that of the car was initialized at the far left (x-axis) and center (of the y-axis). We then moved both of these objects by a specific amount of pixels/s based on a random selection of speeds of these objects. The ball moved vertically downwards, and the car moved horizontally to the right. If the pixels of the ball and car coincided in any frame, that determined a collision, and the ball deflected from its path.

#### Generation of appearance-free videos

To generate a noise pattern consistent with the motion, we first generated a random frame, where p refers to the pixel space. We moved this noise field forward using a flow field *v*(*p, t*) derived from the original video *I*(*p, t*) with an optical flow estimation algorithm, where t refers to the total number of frames in the video. Since *v*(*p, t*) is not an integer field but can contain subpixel motion, the resulting warped results do not perfectly align with the underlying pixel grid. We, therefore, used the nearest neighbor interpolation method to align the newly transformed image to the underlying pixel grid. For more details, refer to Illic et al., 2022^37^. Similar other methods^70^ have also been developed.

### Behavioral Tasks

Videos were presented for various tasks (see descriptions below) on a 24-inch LCD monitor (1,920 × 1,080 at 60 Hz) positioned 42.5 cm in front of the animal.

#### Passive fixation task

During the passive viewing task, monkeys fixated on a white circle (0.2°) for 300 ms to initiate a trial. We then presented a video (300 ms or 600 ms depending on the tasks; see below) followed by a 100 ms gray (background) blank screen, a fluid reward, and an inter-trial interval of 500 ms, followed by the next sequence. We aborted the trials if the gaze was not held within ±2° of the central fixation circle at any point during the video presentation.

#### Binary object discrimination task

Monkeys were trained to fixate on a cross (0.2°) for 300 ms to initiate a trial. The trial began with presenting one image from a pool of 200 for 100 ms. This was followed by a 100-ms blank gray screen. Then, a screen with two images was shown: one was a canonical image of the correct (target) category, and the other was a canonical image of the distractor. The monkeys could look at these images for up to 1500 ms. They indicated their choice by making a saccade and holding their gaze on the target image for 700 ms. If the monkeys did not hold their gaze within a small window (±2°) before the choice screen appeared, the trial was aborted. To facilitate training and behavioral data collection, the monkeys also performed the same task in their home cages ^71^, with the only difference being that they indicated their choice by touching a touchscreen instead of maintaining their gaze on the selected target for 700 ms.

#### Object motion direction estimation task

Monkeys fixated on a cross (0.2°) for 300 ms to initiate a trial. We then presented a video of an object randomly picked from a pool of 200, moving toward one of the eight directions (left, right, up, down, up and left, up and right, down and left, down and right). Then, a blank gray (background) screen was presented for 100 ms, followed by a choice screen with eight small dots (0.2°) located in eight locations corresponding to the eight target movement directions, presented for 300 ms. Monkeys were trained to make a saccade to one of these eight fixed locations based on the perceived direction of motion of the objects. We aborted the trials if the gaze was not held within ±2° of the central fixation circle at any point during the video presentation.

#### Speed discrimination task

Monkeys were trained to fixate on a cross for 100 ms to initiate a trial. A total of 100 videos were presented in the study, each lasting 300 ms and featuring two objects randomly selected (out of a pool of 10), moving with differing speeds – one moving faster (Target) than the other (Distractor) and each presented in a different hemifield. Following each video, a blank screen was presented for 100 ms. After this, canonical images of both objects appeared, and the monkeys had to select the one that had moved faster by making a saccade to that object and fixating on it for 700 ms. The videos contained movement in eight uniformly sampled directions, including four along the cardinal axes and four along the oblique axes.

#### Future event prediction task

We tested the animals on three phases of the task. In phase 1, we showed the animals a training set of 400 videos (full version; 700 ms), while they passively fixated on these videos. The main purpose was to make them aware of the possibilities in these videos (i.e., familiarization with conditions 1-3). Next, on this subset of 400 videos (training set), we trained the animals to perform the future event detection task. Lastly, we used a held-out video (test) set with 105 videos to measure their behavioral reports. Monkeys were shown the first 300 ms of these 105 videos (of a ball and a car in motion, each moving at different speeds). The ball’s movement was directed vertically downward, while the car’s trajectory was horizontal, moving to the right. Depending on their respective speeds, three scenarios were possible: a collision between the two objects (condition 1), the ball passing ahead of the car (condition 2), or the car passing ahead of the ball (condition 3). The speed of the car and ball were selected randomly from the following set: [7 deg/s to 40 deg/s]. The length of the videos was intentionally kept shorter than the entire duration of the event. On the choice screen lasting for 1500 ms, the monkeys were presented with two images, one representing the correct likely scenario and the other one representing one of the two unlikely scenarios. Based on the short video they watched, they had to determine and select the most probable outcome depicted in the video by making a saccade to the image representing the correct scenario and holding their gaze on it for 700 ms (or touching the correct image in the case of the in-cage experiment).

### Neural Recordings

We surgically implanted each monkey with a headpost under aseptic conditions before training on the behavioral tasks. After behavioral training, we recorded neural activity using 10 × 10 microelectrode arrays (Utah arrays, Blackrock Microsystems). A total of 96 electrodes were connected per array. Each electrode was 1.5 mm long, and the distance between adjacent electrodes was 400 μm. Before recording, we implanted each monkey with multiple Utah arrays in the IT cortex and the V4 cortex. IT arrays were placed inferior to the superior temporal sulcus and anterior to the posterior middle temporal sulcus. During each recording session, band-pass filtered (0.1 Hz to 10 kHz) neural activity was recorded continuously at a sampling rate of 20 kHz using Intan Recording Controller (Intan Technologies, LLC). The majority of the data presented here were based on multiunit activity. We detected the multiunit spikes after the raw data was collected. A multiunit spike event was defined as the threshold crossing when voltage (falling edge) deviated by more than three times the standard deviation of the raw voltage values.

### Neural site inclusion criteria

<u>Image rank-order response *reliability per neural site*</u>: To estimate the reliability of the responses per site, we computed a Spearman-Brown corrected, split half (trial-based) correlation between the rank order of the image responses (all images). All neural sites which showed a response reliability greater than zero was used in the analyses.

### Microstimulation of the IT cortex

We used the iridium oxide-coated Utah arrays to perform the microstimulation of the IT cortex. Before the microstimulation experiments, we first selected the electrodes to stimulate. First, we identified all electrodes with an impedance between 100k ohm and 1M ohm. Then, we randomly selected 16 out of these electrodes. We applied 10uA of bipolar pulses for 150 ms starting 50 ms post-video onset. The stimulation pulses were biphasic, with the cathodal pulse leading. Each pulse was 0.2 ms in duration, with 0.1 ms between the cathodal and anodal phases.

### Neural Data Analysis and Statistics

#### Estimating Object Position

We estimated object positions from the neural population activity using cross-validated linear regression (PLS regression with 20 components). The neural responses were first averaged between 70 and 170 ms (post-image onset, similar to ^25^). We then used a 10-fold cross-validation scheme to divide the neural responses and the corresponding x and y object positions (per image) into train and test sets. Then, we estimated the weights and biases for the regression model using the training dataset and tested on the held-out image set. We used a total of 640 images (8 objects, 80 images per object). The goodness of the model was assessed as the Pearson correlation between the neural predictions and the ground truth positions for the x and y positions, respectively.

#### Estimation of object motion direction

##### LSTM classification-based estimates

The neural population responses averaged for 30 ms time bins (from 0 ms to 540 ms) was entirely passed on to an LSTM network (using the bilstmLayer function in MATLAB) with the following parameters: outputSize of 200, and outputMode: ‘sequence’. The output of the LSTM network was then fed to a fullyConnectedLayer (with eight nodes), a softmaxLayer, and a classificationLayer (with eight nodes corresponding to the eight directions used). We used 10-fold cross-validation. For the object identity estimates, we used the same approach, but instead of 8 nodes, we used ten output nodes corresponding to the ten object categories.

##### Linear classification-based estimates

For the linear classification (**Figure 2E-F, 3E-F**), the neural population responses averaged for 30 ms time bins (from 0 ms to 570 ms) were used independently with linear classifiers (linear discriminant analysis). We used 10-fold cross-validation across the videos to train eight one-vs-all LDA classifiers, each representing one specific motion direction.

#### Estimation of object speed

##### LSTM regression-based estimates

The neural population responses averaged for 30 ms time bins (starting from 0 ms to 570 ms) were entirely passed on to an LSTM network (using the bilstmLayer function in *MATLAB*) with the following parameters: outputSize of 200, and outputMode: ‘sequence’. The output of the LSTM network was then fed to a fullyConnectedLayer (with one node), and a regressionnLayer (with one node corresponding to the output speed). We used 10-fold cross-validation.

##### Linear regression-based estimates

For the linear regressions (in **Figure 5B**), the neural population responses averaged for 30 ms time bins (from start times of 0 ms to 570 ms) were used independently with linear regression (PLS regression). We used 10-fold cross-validation across the videos to train the PLS regression parameters (with 20 components) for speed prediction.

#### Predicting the speed discrimination task accuracies

We averaged the neural responses per site according to various options of *tStart* and *binwidth* (inset, **Figure 6C**) to generate multiple sets of population responses per video. Each parameter setting of *tStart* and *binwidth* can be considered as a specific decoding algorithm (indicated as separate dots in **Figure 6C**). For each choice of decoding algorithm, we then regressed the population responses onto the video-by-video task accuracy estimated from the behavioral measurements. This was done using a 10-fold cross-validation. Therefore, we generated held-out predictions for each video (when it was part of the held-out test set). The predictions were correlated (Spearman correlation) with the empirical behavioral accuracies to estimate the consistency between the neural decodes and the behavior. We also estimated the trial split-half correlation (internal consistency) for the video-by-video behavioral accuracies and the neural predictions (per decoder). The square of the multiplication of these two estimates served as the noise ceiling for each decoder. The values in the y-axis of **Figure 6C** and **D** are the Spearman correlations corrected (divided by) the individual noise ceilings.

#### Predicting the future events from neural data

Each video belonged to one of the three outcomes, based on whether the ball and the car collided. Depending on their speeds, they would either collide (*condition 1*), the ball would move past the car (*condition 2*), or the car would move past the ball (*condition 3*). We trained a linear classifier (regularized LDA) to map the neural data (population responses) averaged across specific time intervals post video onset (as described in earlier analyses) to each video label in a cross-validated way (10-fold cross-validation). The IT-based video-by-video average accuracies were estimated on held-out sets and correlated (Spearman R) with the measured video-by-video average behavioral accuracies (as shown in **Figure 8D**).

### Statistical analyses and significance testing

For most analyses (as mentioned specifically in the Results text), standard parametric paired t-tests and Pearson and Spearman correlations were used. For other cases, we used non-parametric permutation-based null distribution estimation and statistical tests (see below).

*Permutation tests to assess the statistical significance of decoder performances*.

We used a permutation test to estimate the statistical significance of decoding accuracy for motion speed or direction for the above-mentioned decoding approaches. We randomly shuffled the direction, speed, or event condition labels (repeated 1000 times) and re-performed the decoding analysis for each repeat to construct a null hypotheses space. A timepoint was declared statistically significant if the values with the non-shuffled decoding accuracy were above the 95% CI (i.e., the standard deviation) of the null distribution.

## Acknowledgments

KK has been supported by funds from the Canada Foundation for Innovation (CFI), the Canada Research Chair Program, the Simons Foundation Autism Research Initiative (SFARI, 967073), and the Canada First Research Excellence Funds (VISTA Program). HR is funded by CIHR Postdoctoral Fellowship. FI was supported by the Canada First Research Excellence Fund (VISTA Program). RPW is supported by the Canada First Research Excellence Fund (VISTA Program) and the National Sciences and Engineering Research Council of Canada (NSERC). We thank Jim DiCarlo (JJD) and the members of the DiCarlo Lab for their help with data collection and curation resources supported by the Simons Foundation (542965, JJD).

## Supplementary Material

**Figure S1.**
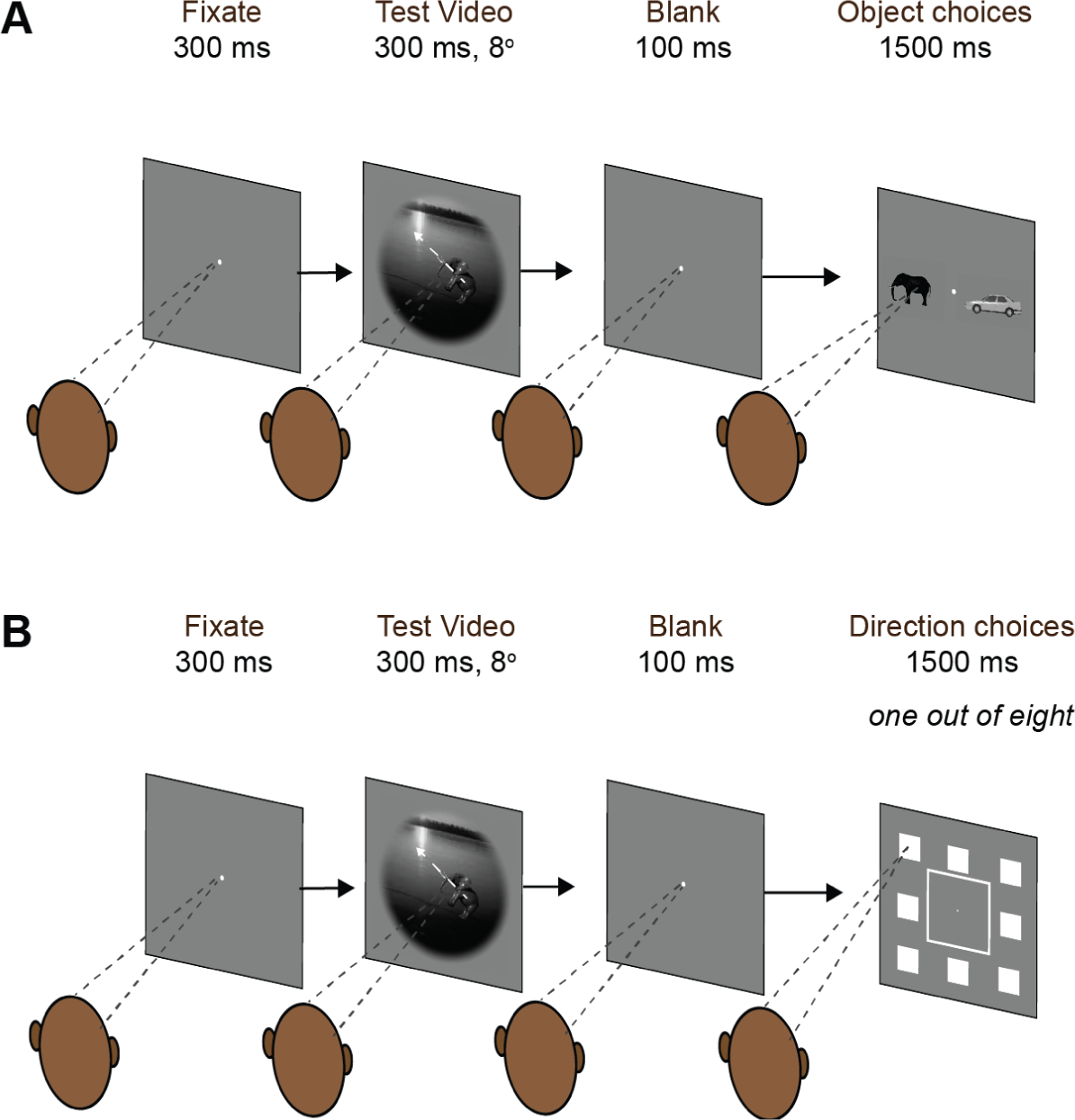
A. Object identity recognition task. Monkeys fixated on a small white dot for 300 ms, after which a Test video is presented for 300 ms, showing an object moving within an 8° background. After a brief 100 ms blank screen, we presented canonical images of those two objects for 1500 ms, and the monkeys had to select the one that matched the object identity during the Test video by making a saccade to that object. **B.** Object motion direction discrimination task. Similar to the object identity task in A, except that in the response phase monkeys were tasked with indicating the motion direction of the object by making a saccade towards one of the eight possible locations corresponding to the motion direction, within 1500 ms.

**Figure S2.**
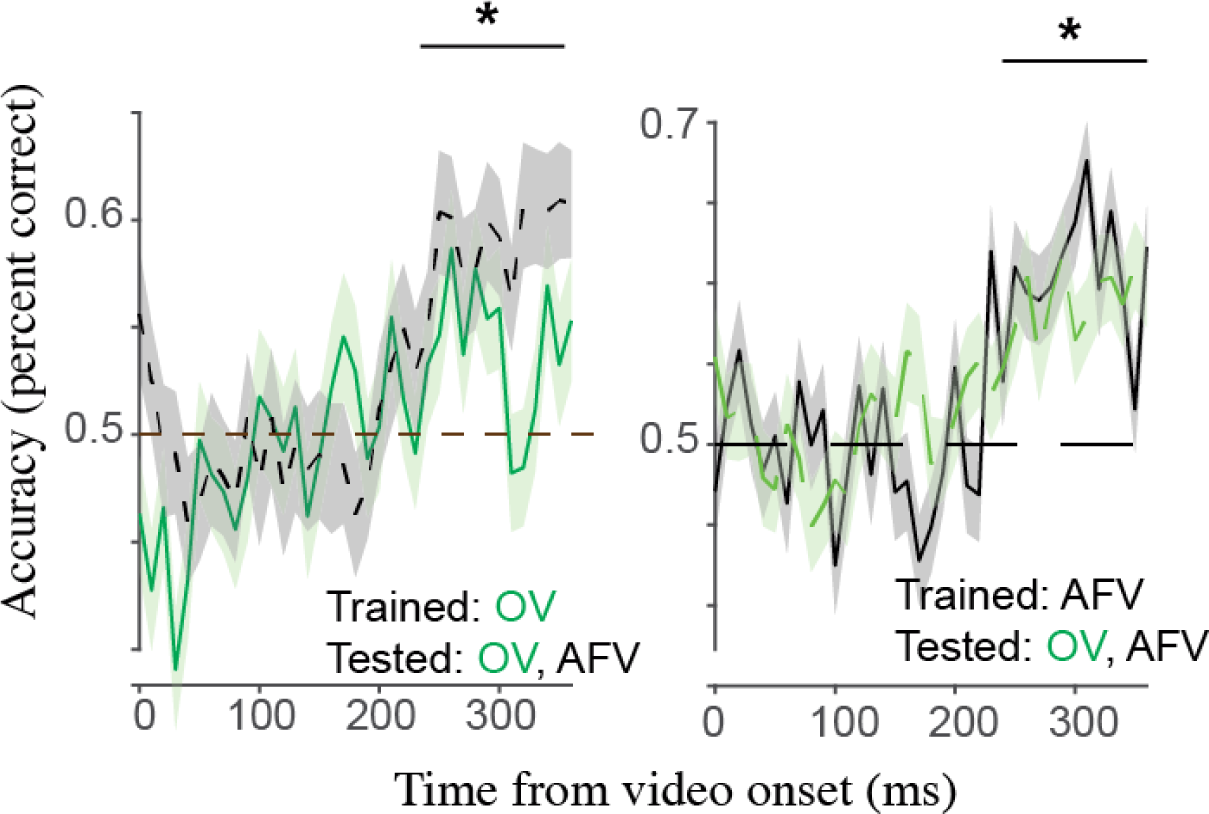
Decoding motion directions from IT population activity for videos with and without appearance information. LSTM-based decoding models linking IT population activity to object motion direction discrimination perform well above chance levels (p<0.05, permutation test) approximately 200 ms post video onset for both appearance-free videos (AFV) and original videos (OV), with the left graph representing models trained on OV and tested on OV (green) and AFV (black), and the right graph depicting models trained on AFV and tested on both OV (green) and AFV (black). In both cases, object motion direction decoding accuracy reaches a level significantly above the chance level, as indicated by asterisks in the figure. Errorbar bands denote s.e.m across videos.

**Figure S3.**
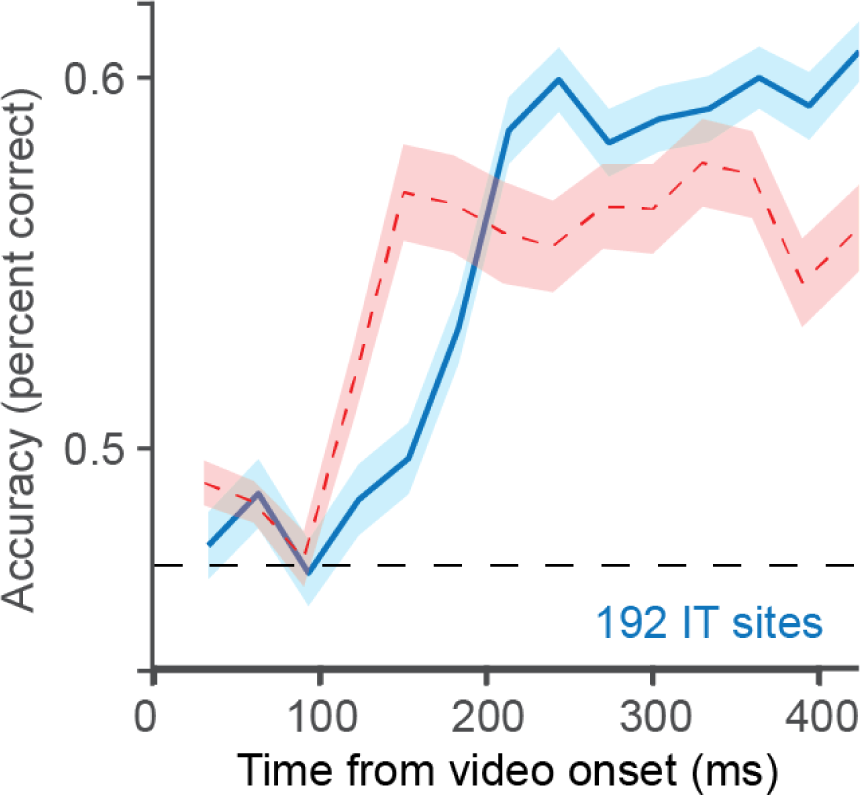
Comparison of dynamics of object motion identity and speed decodes from the IT population activity. Both object identity (in red) and motion speed (in blue) have been estimated using a regularized LDA classifier based on the instantaneous (30 ms averaged) population responses across 192 IT sites (pooled across two monkeys). Similar to motion direction decodes, speed estimates lag behind the object identity estimates.

